# DEVELOPMENT OF A BIOMIMETIC 3D OVARIAN SCAFFOLD USING DECELLULARIZED EXTRACELLULAR MATRIX AND MECHANICALLY TUNED HYDROGELS

**DOI:** 10.64898/2026.03.07.709996

**Authors:** S.K Nikhila, Saran S Menon, V M Vanishree, Guruprasad Kalthur, Sneha Guruprasad Kalthur, Ankitha Suresh, Bhisham Narayan Singh, Kirthanashri Vasanathan, N.S Raviraja, Ramya Nair

## Abstract

Artificial ovarian scaffolds represent a promising therapeutic strategy for preserving reproductive health in patients. However, current *in vitro* approaches are limited by inadequate biomimicry of the native tissue microenvironment, leading to poor development of *in vitro* ovarian models. In this study, we developed region-specific hydrogel scaffolds incorporating solubilized decellularized ovarian extracellular matrix (dECM) with mechanically tuned properties to enhance the functionality of engineered 3D ovarian models. Ovine ovarian dECM was isolated by mechanical and chemical decellularization methods and subsequently solubilized and incorporated in varying concentrations in homogenous alginate (0.5%) and a composite mixture of 1% gelatin with 0.5% alginate (1:1). The synthesized hydrogels were characterized for rheological properties, including Young’s modulus, pore size, and viscosity, and cytocompatibility assays were conducted using Chinese hamster ovary (CHO) cells. The study demonstrated that both 0.5% alginate and the composite gelatin-alginate hydrogels successfully replicated the mechanical properties of native human ovarian cortical and medullary tissue, with Young’s modulus of 0.84 ± 0.16 kPa, pore size (60-150 nm), and toughness of 0.4Pa, respectively. Zonal hydrogel scaffolds incorporating ovarian dECM demonstrated significantly enhanced cell viability compared to hydrogels supplemented with dECM. The study emphasises the critical role of integrating both mechanical and biochemical attributes while developing functional artificial ovarian constructs for transplantation and regenerative medicine applications. This work contributes to advancing strategies for creating physiologically relevant *in vitro* models of ovarian tissue.

## Introduction

The ovary functions as a finite reservoir for germ cells and as a dynamic endocrine organ essential for female reproductive and systemic health. Disruptions in ovarian function can arise from environmental factors(1), metabolic(2) and autoimmune disorders(3), viral infections(4), and genetic predisposition(5) potentially leading to infertility and long-term health complications, including osteoporosis, cardiovascular disease, autoimmune disorders, and depression(6). Furthermore, the rising survival rates among young cancer patients highlight the need to address fertility concerns, as gonadotoxic radiotherapy and chemotherapy have been associated with decreased primordial follicle count, increased vascular damage, and ovarian cortical fibrosis(7). Infertility is a growing global issue, affecting 17.5% adult population (1 in 6) according to the World Health Organization (WHO) 2023, highlighting the magnitude. Hence, addressing fertility concerns is becoming imperative with changes in environmental conditions, the lifestyles of the younger population, and advances in treatment options.

Fertility preservation is a practical approach to safeguarding reproductive health, considering environmental, medical, and age-related factors. Current medical strategies for addressing ovarian dysfunction include ovarian and uterine transposition, vitrification of embryos and oocytes, and ovarian cortex cryopreservation(8,9). A notable aspect of tissue cryopreservation is its ability to utilize either fragments or entire ovaries for autotransplantation, thereby circumventing the need for immunosuppression and eliminating the concern of organ rejection. Despite their clinical utility, these methods pose a significant limitation: the potential risk of reintroducing malignancy(10,11). In this context, bioengineered ovary reconstruction represents a promising frontier.

Advances in biomedical engineering have led to the creation of artificial ovarian structures that can support the maturing follicle survival and endocrine function(12). Various hydrogels and microfluidic cultures report the successful development of follicles, however, the native ovarian heterogeneity and architecture are not reproduced in these conditions(13,14). Reproducing the native ovarian extracellular matrix (ECM) proteins within polymers poses a notable challenge, as these proteins are essential for activating signalling molecules that play a crucial role in ovarian function(15). Decellularized ovarian tissue provides this natural bio-matrix, providing a supportive environment that can facilitate the restoration and regulation of ovarian functions(16). However, no current model simultaneously recapitulates both the biochemical and biomechanical heterogeneity of the human ovary, limiting translational relevance.

Despite these advances, current *in vitro* ovarian models fail to consistently support human folliculogenesis, primarily because of the absence of physiologically relevant biomechanical cues and spatial heterogeneity(17). In addition to biochemical factors, research has demonstrated variations in the mechanical properties of the ovary, including stiffness, porosity, and shear stress, across reproductive and nonreproductive stages(18). Studies have also proven the role of the softer medullary layer in the ovary in supporting the growth and expansion of secondary and hormone-producing antral follicles(19). These findings underscore mechanotransduction as a critical yet underexplored regulator of folliculogenesis and endocrine function, operating alongside biochemical signalling.

In this study, we hypothesize that integrating solubilized decellularized ovarian extracellular matrix (dECM) with region-specific mechanically tuned hydrogels will enhance the functionality of the 3D ovarian model *in vitro*. This approach represents a crucial step in developing a 3D model of the human ovary, with potential applications not only as a transplant to improve fertility preservation but also as a model for *in vitro* drug toxicity analysis. Beyond fertility preservation, this platform offers a physiologically relevant *in vitro* model for studying ovarian biology, mechanobiology, and reproductive toxicology. Collectively, this work advances the field of ovarian tissue engineering by bridging the gap between biochemical composition and mechanotransduction, paving the way for clinically translatable artificial ovary platforms.

## Materials and methods

### Ovary collection

Ovine ovaries were obtained from a local slaughter house and transported within 3 h timeframe to the laboratory in phosphate-buffered saline (PBS) solution containing 1% antibiotics (IAEC/KMC/15/2023). Upon arrival, the ovaries were meticulously separated and thoroughly cleaned in DMEM medium (Cat. No. 12800017, Gibco, India) supplemented with 5% penicillin-streptomycin (Cat. No. 15140-122, Gibco, India). The ovaries were horizontally cut into 2 mm-thick sections. The sectioned ovaries were randomly assigned to either the control group or the decellularized group. Ovaries in the control group underwent RNA isolation and fixation in paraformaldehyde (Cat. No: 30525-89-4, Merck, Germany), while the remaining ovaries were used for decellularization.

### Decellularization of ovaries

The ovarian tissues were decellularized by mechanical and chemical methods as detailed by Hosseinpour et al.,(20) with minor modifications. Briefly, ovarian sections stored at -80°C were incubated at 37°C for 30 min, and this process was repeated thrice. Subsequently, the sections were stirred at 150 rpm for 3 h in a solution of 0.5% (w/v) sodium dodecyl sulfate (Cat: 151-21-3, Sisco Research Laboratory Pvt.Ltd., India), followed by a triple rinse in distilled water and an overnight agitation in 1% (v/v) Tween 20 (Cat: 9005-64-5, Central Drug House (P) Ltd., India). The sections were subjected to hypotonic treatment (deionized water) for 9 h, followed by a detergent wash using 2% (w/v) sodium deoxycholate (Cat. No. 302-95-4, Tokyo Chemical Industry Pvt. Ltd., India) for 12 h. The ECM underwent an extensive cleansing process, including 6 h of washing with MilliQ water to ensure complete removal of detergents. The water was replaced every 2 h. The decellularized sections were assessed for histological and DNA analyses to assess the quality of the ECM following the decellularization procedure(21), and the remaining sections were used for solubilization.

### Solubilization of the ovary

Decellularized ovarian sections were digested using 60 mg of pepsin in hydrochloric acid at pH2, maintained at 37°C for 24 h. Once the ovarian sections were completely dissolved, the solution pH was restored to 7.2 with 10 N NaOH (Cat. No: 1310-73-2, Sisco Research Laboratory, India), and the solution was maintained at 4°C. This solution was lyophilized overnight, and the resulting powder was analyzed for protein quantification and toxicity(22).

### DNA analysis

The native and decellularized sections, each weighing 100 mg, were homogenized (Cat. No: BT704, Benchtop) and solubilized in 200 µL of Radioimmunoprecipitation Assay buffer (RIPA; Cat. No: TCL131, Himedia Laboratories, India) at 56°C. Following this, 200 μL of Trizol (Cat. No: 9108, DSS Takara Bio India Pvt. Ltd.) was added to the homogenised samples, and incubated for 5 min at room temperature and centrifuged at 12,000 g for 15 min at 4°C. In the next step, chloroform was added, and DNA was precipitated from the aqueous phase by adding an equal volume of isopropanol (Cat.No: 67-63-0, Sisco Research Laboratory Pvt.Ltd., India) followed by an incubation period of 1 h at -20°C. Centrifugation was performed to collect the pellet, which was washed with 100% ethanol and then 70% ethanol (Cat. No: 64-17-5, American Chemical Society, Washington, D.C.). The resultant pellet was dissolved in RNase-free water (10 μL), and the DNA concentration was determined using a Nanodrop (SHIMADZU Biotech) by analysing the absorbance at 260/280 nm. The amount of DNA was averaged across three independent runs and expressed as ng/mL (23).

### Protein analysis

Native and decellularized tissue samples were homogenized in 200 µL of RIPA buffer containing a protease inhibitor (Cat. No: A32961, Thermo Fisher, India). The homogenates were centrifuged at 12,000 × g for 10 min at 4 °C, and the supernatant was collected for protein estimation(24).

Protein concentration was determined using the Bicinchoninic Acid (BCA; Cat. No: 23228, ThermoFisher Scientific Inc., India) assay according to the manufacturer’s protocol. Briefly, BCA working reagent was prepared by mixing reagent A and reagent B (50:1). 25 µL of sample were added to a 96-well plate, followed by 200 µL of BCA working reagent. The plate was incubated at 37 °C for 30 min, and absorbance was measured at 562 nm using a microplate reader(24).

### Scanning Electron Microscopy

The lyophilized ovarian tissues, both decellularized and non-decellularized, were subjected to serial graded ethanol-water solutions (30, 50, 70, 80, 90 and 100 % ethanol) for 15 min each after fixation with 2.5% glutaraldehyde overnight. The fixed samples were coated with a gold layer and imaged using an ultra-high-resolution scanning electron microscope ( EVO MA18 with Oxford EDS(X-act))(25).

### Histological analysis

For histological analysis, native and decellularized tissue samples were fixed in 4% PFA (Cat. No: 30525-89-4, Merck, Germany) at room temperature. The samples were dehydrated in graded alcohol (70%, 80%, 90%, 95%, and 100%), cleared with xylene (Cat.No: 1330-20-7, Sisco Research Laboratory Pvt.Ltd., India), and embedded in paraffin. After dewaxing and rehydration, serial microtome sections (5 µm) were stained with hematoxylin and eosin (H&E) and 4 µg/mL of 4’,6-diamidino-2-phenylindole (DAPI; (Cat. No: D1306, ThermoFisher Scientific Inc., India) to confirm the absence of nuclear material in the decellularized sections(26).

### Hydrogel preparation

Sodium alginate and gelatin–sodium alginate hydrogels were prepared with varying polymer compositions: sodium alginate alone (0.25, 0.5, and 0.75% w/v) and in combination with gelatin (0.5, 1, and 2%) and 0.5% alginate (1:1 ratio). Sodium alginate (Cat. No: 9005-38-3, Sisco Research Laboratory Pvt. Ltd., India) solution was prepared by dissolving the alginate powder in Milli-Q water for 2h at 50℃ (27). The prepared sodium alginate solution was crosslinked with 1% Calcium chloride (CaCl₂) (Cat. No. 10043-52-4, Merck) for 2 min to form a gel. Separately, 1% (w/v) gelatin (Cat.No: 9000-70-8, Sisco Research Laboratory Pvt.Ltd., India) & 0.5% (w/v) sodium alginate composite hydrogel was prepared by mixing gelatin and sodium alginate in milliQ water under constant stirring at 50 rpm for 1hr at 50°C, followed by the addition of crosslinker EDC/NHS (2:1) for 2–3 h at 100 rpm at 50°C(28). The hydrogel mixture thickened and was crosslinked with 1% CaCl₂ to obtain stable hydrogels.

For extracellular matrix (ECM) incorporation, lyophilized decellularized ECM (dECM) of different concentrations (1, 2.5, 5, 50, 100, 500, and 1000 µg/ml) was supplemented to the hydrogels before solidification.

### Degradation and Swelling

The degradation rate of the hydrogels was evaluated by immersing the hydrogels in DMEM culture medium maintained at 37°C. The dry weight of each hydrogel was measured for 5 min every half hour, followed by a 24 h gap. Their water-intake capacity was evaluated by measuring the swelling rate of lyophilised hydrogels incubated in DMEM at 37°C to maintain physiological conditions. At each time point (15, 30, 45, 60, and 120 min), the dry weight of the hydrogels was calculated until there was a decrease in weight (29).

### Brunauer–Emmett–Teller (BET) analysis

To understand the surface characteristics and porosity of the prepared hydrogels, Brunauer–Emmett–Teller (BET) analysis was performed in lyophilised hydrogels. Nitrogen adsorption–desorption isotherms were recorded at 77 K using a (MICROTRAC – BELSORP MINI X) analyzer. The surface area and the pore size distribution of hydrogels was determined using the Barrett-Joyner-Halenda (BJH) method (30).

### Universal Testing Machine (UTM) analysis

The stiffness and elasticity properties of the hydrogels were tested using a universal testing machine (Instron 3369, USA). The hydrogel doublet samples (20 mm × 20 mm × 5 mm ± 1) were tested in uniaxial compression (or tension) at a crosshead speed of 1 mm/min. Load and displacement were recorded and converted to engineering stroke force to obtain the hardness/compression graph(31).

### Viscosity

The rheological properties and viscosity of the hydrogel formulations were analyzed using a rotational rheometer (Anton Paar MCR 302, Austria). Measurements of doublet samples were performed in a parallel-plate geometry (25 mm) with a 1 mm gap, maintained at 37 °C to simulate physiological conditions.

The flow behaviour of the hydrogels was assessed by analysing steady-state shear tests over a range of 0.1–100 s⁻¹. The viscosity profile was plotted against the shear rate to evaluate the shear-thinning characteristics. Oscillatory frequency sweep tests were performed in the range of 0.1–100 Hz within the linear viscoelastic region(29).

### Cell culture

Chinese hamster ovary (CHO) cell lines were purchased from NCCS, Pune. The cells were cultured in DMEM with 10% (v/v) FBS (Cat. No. A5256701, Gibco, USA) and 1% (v/v) penstrep (Cat. No. 15140-122, Gibco, India) and maintained at 37 °C in a humidified 5% CO_2_ atmosphere. Once the cells reached 80% confluence, they were mixed along with the hydrogel solution and then crosslinked to form hydrogel beads. The hydrogel beads were further cultured *in vitro*(32).

### 3-(4,5-dimethylthiazol-2-yl)-2,5-diphenyltetrazolium bromide (MTT) assay

Cell viability was assessed using the 3-(4,5-dimethylthiazol-2-yl)-2,5-diphenyltetrazolium bromide (MTT) assay (Cat. No: M6494, ThermoFisher Scientific Inc., India). The culture medium (Dulbecco’s Modified Eagle Medium) was first removed, and the cells were gently washed twice with Dulbecco’s phosphate-buffered saline (DPBS; Cat. No: 2160010, ThermoFisher Scientific Inc., India) for 5 minutes each. Subsequently, MTT solution (0.5 mg/mL) was added to each well of a 96-well plate at a volume of 100 µL and incubated at 37 °C for 2–3 hours to allow the formation of formazan crystals. After incubation, the MTT solution was carefully removed, and dimethyl sulfoxide (DMSO; Cat. No: 67-68-5, Sisco Research Laboratory Pvt. Ltd., India) was added to solubilize the crystals. The plate was then kept at room temperature in the dark for 30 minutes to ensure complete dissolution (33). Finally, the absorbance was recorded at 570 nm, 590 nm, and 630 nm using a microplate ELISA reader.

### Statistical Analysis

The data were statistically analyzed using GraphPad Prism 9 (GraphPad Software, San Diego, CA, USA). The different groups were compared using analysis of variance (ANOVA), followed by the Tukey test (as a post hoc test). All data are presented as the mean and standard deviation of the mean (mean ± SD). In all analyses, p≤ 0.05 was considered statistically significant.

## Results

### Decellularization of ovine ovaries to obtain decellularized samples

Ovine ovaries (n=3) were selected for the study because of their resemblance to human ovaries in terms of structure and follicle arrangement (34). Before decellularization, ovine ovaries were sliced into strips (Figure 1a-c). The strips were sequentially decellularized, and macroscopic observations revealed that the ovaries maintained their shape and homogeneity without any deformation (Figure 1c,d). The change in colour from pink to white indicated preliminary changes in cellular components (Figure 1c,d) and a decrease in ovarian weight from 3.327 g to 2.163 g. The successful decellularization was confirmed by H&E staining, which showed the removal of cellular remnants and nuclei from dECM, while nuclei were clearly visible in native tissues (Figure 2a).

**Figure 1:**
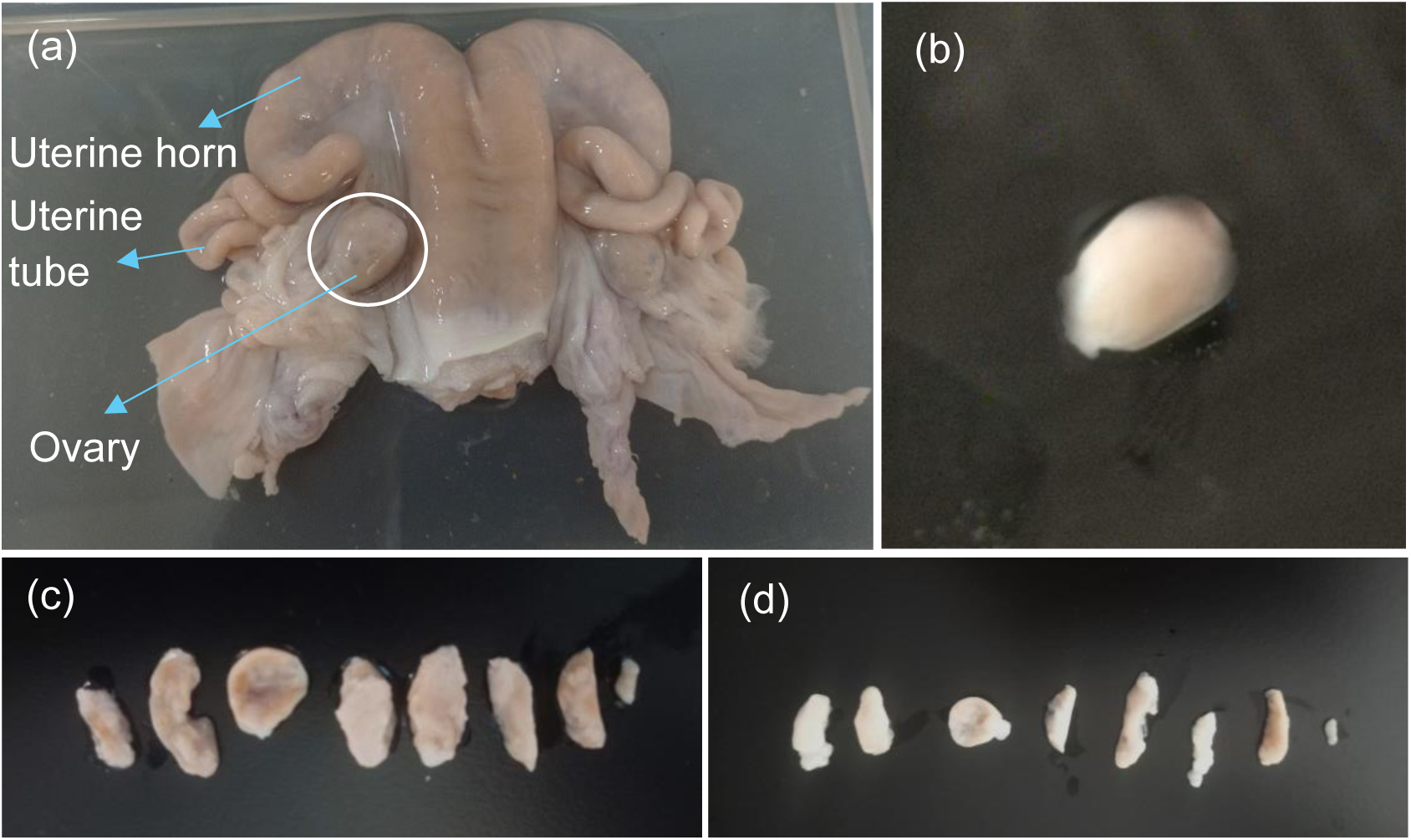
Representative images of (a) female ovine reproductive system, (b) Ovine ovary (c,d) Sectioned ovary before and after decellularization respectively

**Figure 2:**
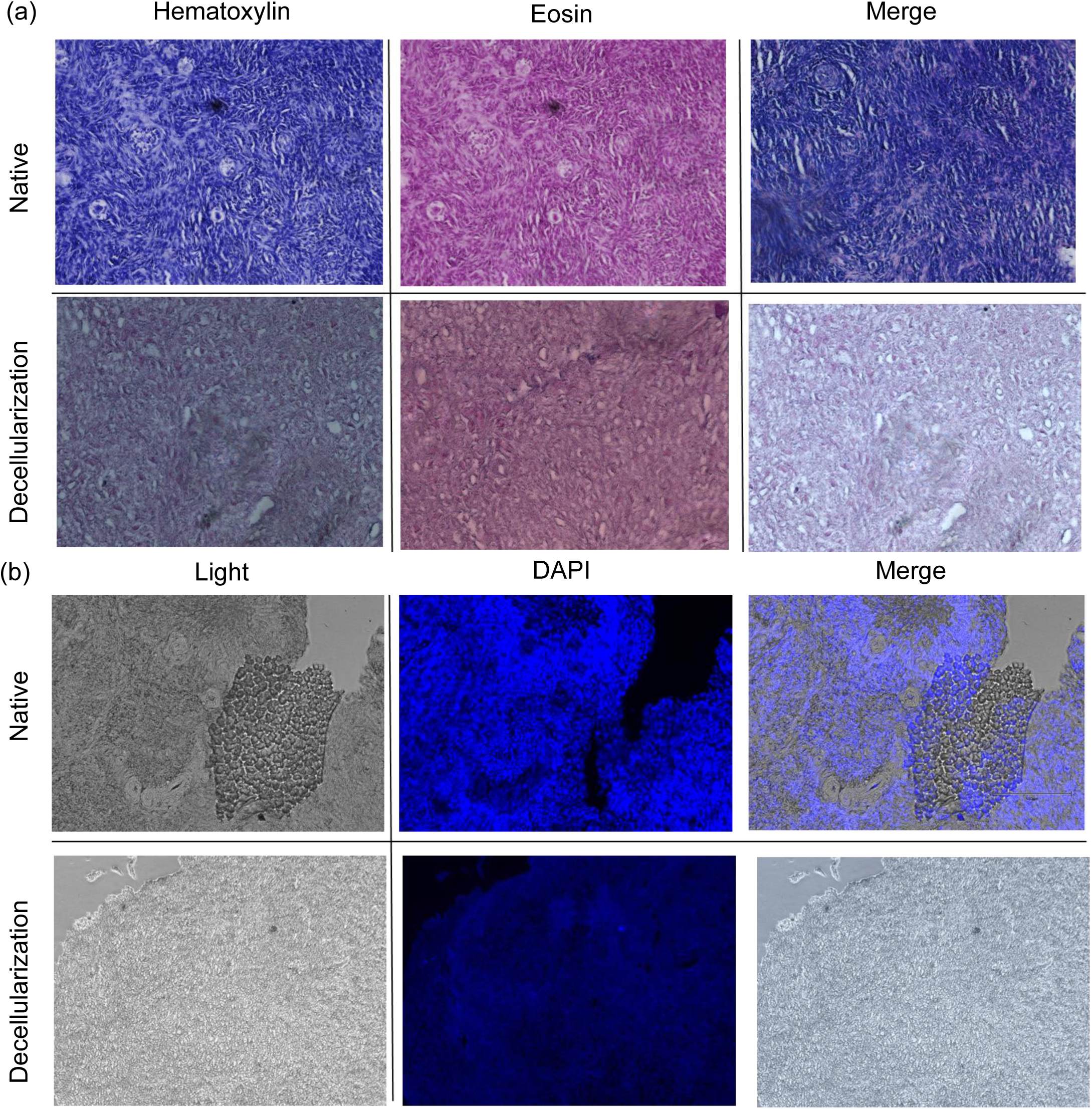
Histological evaluation of decellularized ovary: Representative images of a) H& E staining and b) DAPI stained sections of native and decellularized ovarian sections

In agreement with the histological data, the quantified DNA showed a decrease in DNA content in decellularized ovaries (0.861 ± 0.042 ng/mL) compared to native tissue (1.6055 ± 0.195 ng/mL) (Figure 3b). Successful decellularization was confirmed by scanning electron microscopy (Figure 3a), which revealed cell removal, preserved cell-free cavities, an intact extracellular matrix (ECM) framework, and well-connected, oriented collagen fibers.

**Figure 3:**
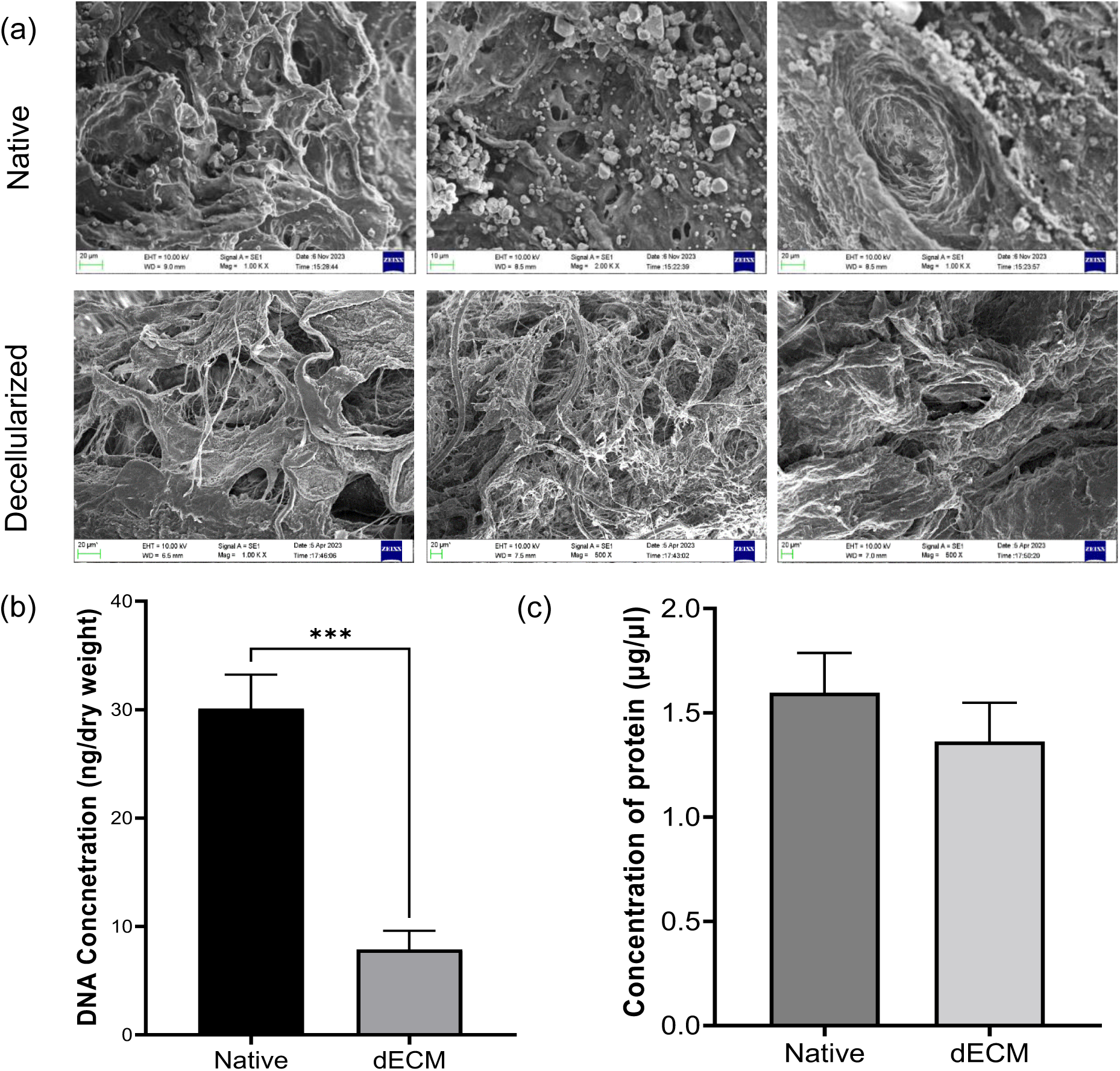
Quality determination of decelluarised extracellular matrix post decellularization: a) Scanning electron microscopy images of native and decellularized ovary, b) Graph representing the DNA content in ovary pre and post decellularization, c) Protein content of native and decellularized ovary analysed by BCA assay

### Protein estimation in decellularized ovary

The total protein content of the decellularized ovary (n=3) was compared with that of the native ovary before and after solubilization (Figure 3c). The decellularized ovarian proteins exhibited a protein concentration of 1.363 ± 0.185 µg/mL, as determined by the BCA assay. This concentration closely approximates that of the native tissue, which was recorded at 1.597 ± 0.185 µg/mL.

### Cytotoxicity of dECM laden hydrogels

The cytocompatibility of hydrogels incorporating varying concentrations of solubilized and lyophilized dECM proteins was evaluated using the MTT assay on CHO cells after 14 days of culture (Figure 4a).

**Figure 4.**
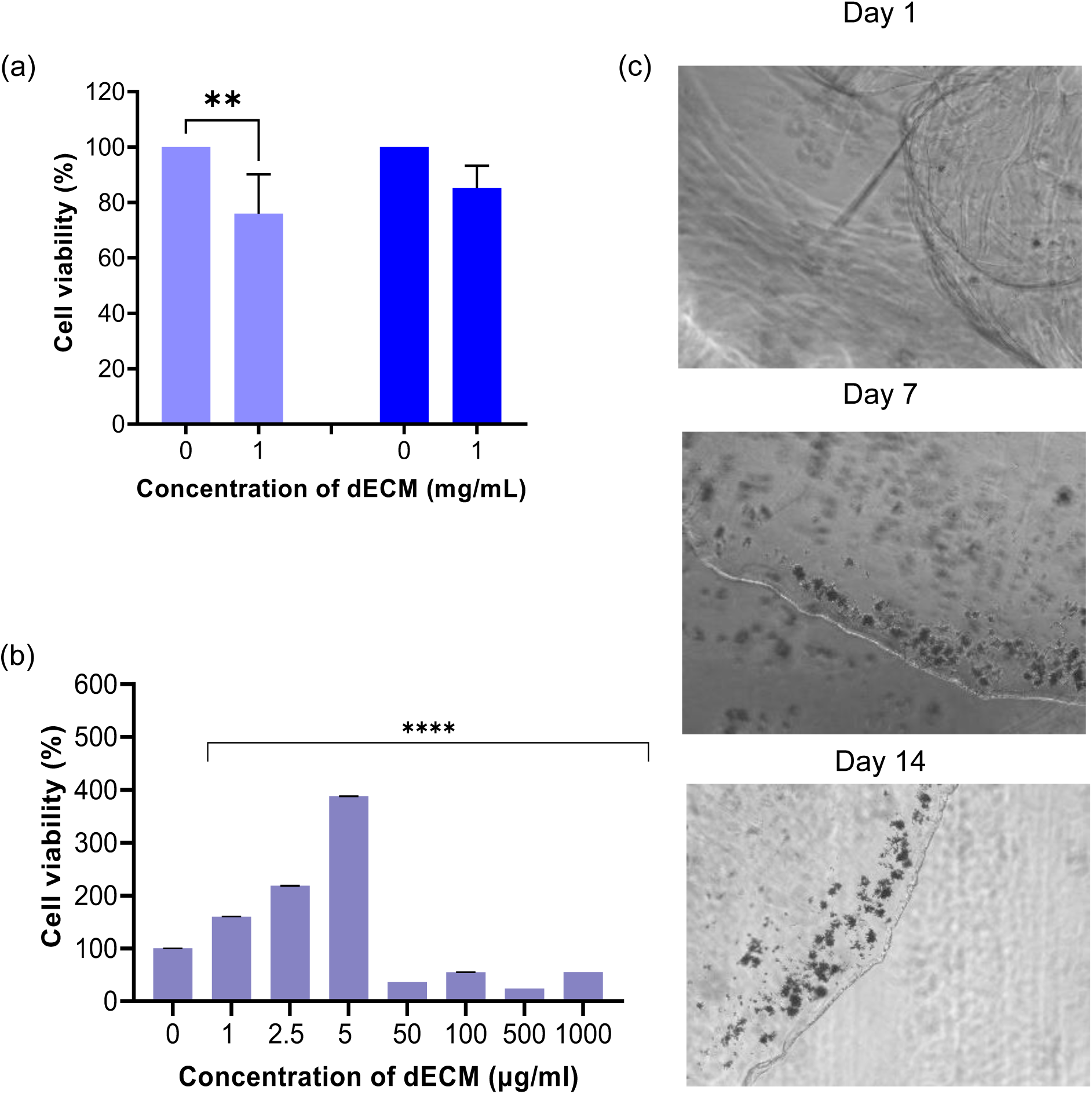
: MTT analysis: a) 1mg/ml of dECM in1% & 2% gelatin with 0.5% sodium alginate (p<0.01 Vs control) [light blue 0-1% gelatin with 0.5% sodium alginate; light blue 1-1% gelatin with 0.5% sodium alginate and dECM; dark blue 0-2% gelatin with 0.5% sodium alginate; dark blue 1-2% gelatin with 0.5% sodium alginate and dECM], b) varying concentration of dECM (p<0.0001 Vs control), with sodium alginate,(c) representative images of the cells cultured in 0.5% of sodium alginate

A significant decrease in cell viability was observed in the 1% gelatin with 0.5 % alginate hydrogel (5) upon supplementation with 1 mg/mL of ECM proteins, yielding a viability of 75.96 ± 14.21% (p < 0.01) (Figure 4a). In contrast, hydrogel containing 2% gelatin with 0.5% alginate exhibited a non-significant decrease in cell viability, measuring 85.14 ± 8.108 %. Moreover, supplementation of 1 mg/mL in 0.5% alginate was found to be highly toxic to CHO cells with a viability of 55.19 ± 0.03% (p<0.0001) the cell toxicity and ECM proteins was observed at all the higher concentrations used, from up to 50 µg/mL (36.21 ± 0.02, p<0.0001) to 1 mg/mL (Figure 4b). Conversely, a dose-dependent increase in cell viability was observed with lower concentrations of ECM protein (1–5 µg/mL; 387.86 ± 0.24 at 5 µg/mL, p<0.0001). For further studies, a 2% gelatin-alginate mixture was not considered due to the difficulty in molding the hydrogel. These findings indicate that both the hydrogel composition and ECM protein concentration regulate cell viability.

### Degradation ratio and Swelling ratio

The swelling rate of the hydrogels was studied by measuring the absorption rate of lyophilized hydrogels immersed in Dulbecco’s modified eagle medium (DMEM), and the deswelling rate was measured by analyzing the change in mass of the hydrogels initially every 15 min for 2 h, followed by every hour until 6 h, and then a 24-h gap until the hydrogel degraded.

Within 15 min of immersion, the maximum swelling rate was observed in both hydrogels containing 1% gelatin, achieving approximately 1.83 ± 0.8 g, while alginate reached approximately 1.38 ± 0.3 g. A stable hydrogel mass was observed for up to 120 min; thereafter, a deswelling phase was observed in the alginate hydrogel at 120 min, indicating an initial phase of mass loss (Figure 5c). In contrast, the combination of gelatin and alginate exhibited a more stable swelling and degradation rate than alginate alone.

**Figure 5:**
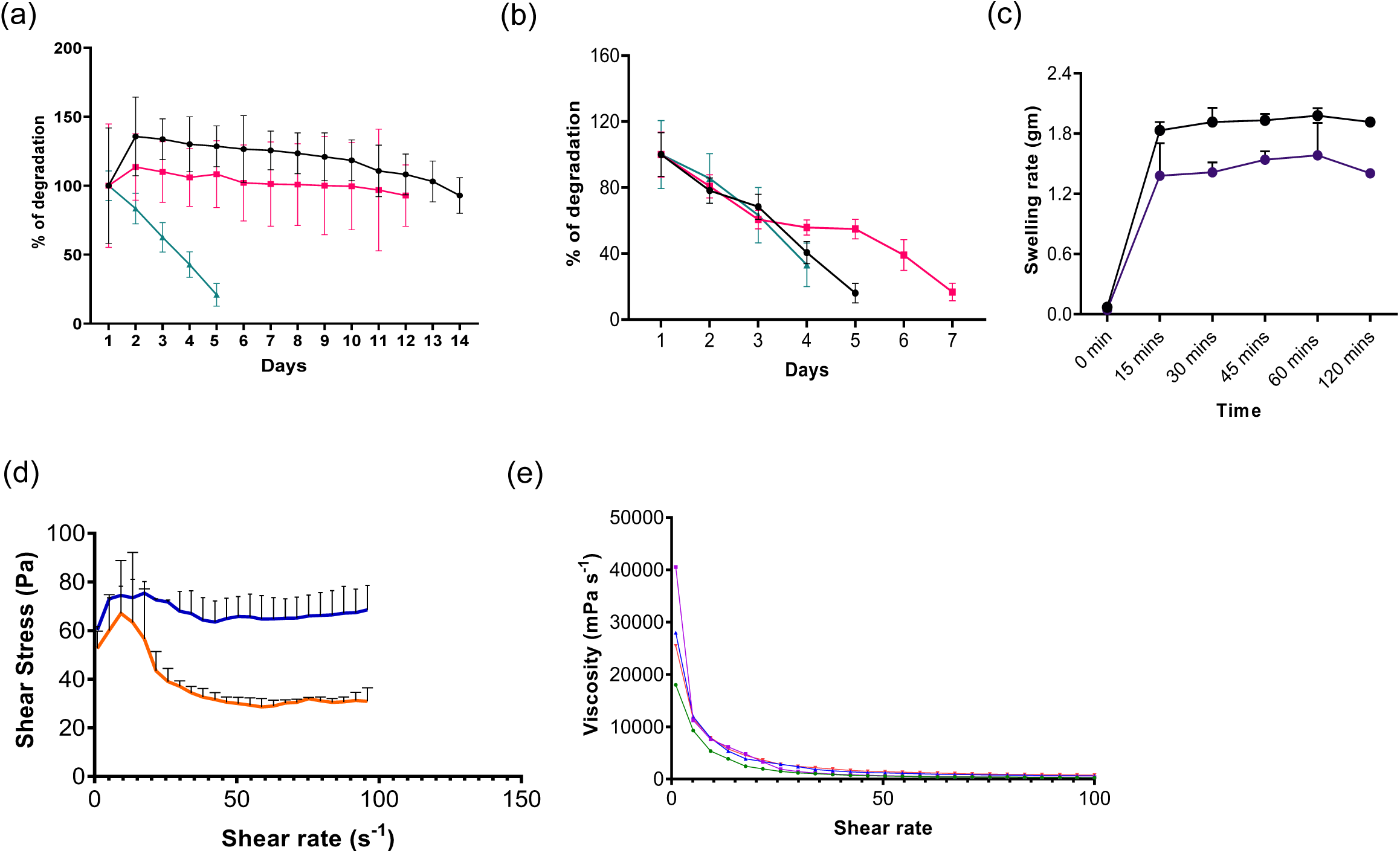
Percentage of degradation of: (a) 0.75%, 0.5% and 0.25% Sodium alginate [black-0.75%, pink-0.5% and blue-0.25%], (b) 2%, 1% & 0.5% gelatin with 0.5% sodium alginate [black-2% gelatin+0.5% sodium alginate, pink-1% gelatin+0.5% sodium alginate and blue-0.5% gelatin+0.5% sodium alginate]; (c) swelling rate of 0.5% Sodium alginate and 1% gelatin with 0.5% sodium alginate; Rheological analysis of the hydrogels (d) Shear stress rate for 0.5% Sodium alginate and 1% gelatin with 0.5% sodium alginate; (e) viscosity-shear rate graph for 0.5% Sodium alginate and 1% gelatin with 0.5% sodium alginate [orange-0.5% sodium alginate; blue-1% gelatin+0.5% sodium alginate]

The degradation rates of homogenous sodium alginate at varying concentrations of 0.25, 0.5, and 0.75% were systematically evaluated. The 0.75% sodium alginate gradually decreased from 19.6 ± 8.19 mg to 18.12 ± 2.52 mg over 14 days, 0.5% sodium alginate decreased 17.01 ± 8 mg to 16. 64 ± 3.9 mg within 12 days and 0.25% had rapid degradation from 18.7 ± 2 mg to 3.9 ± 1.5mg, followed by structural breakage and complete dissolution of the gel (Figure 5 a).

The degradation rate of a composite polymer comprising gelatin combined with 0.5% sodium alginate was assessed. The 2% gelatin with 0.5% sodium alginate and 0.5% of gelatin and 0.5% sodium alginate resulted in rapid degradation by reduction in weight from 28.14 ± 3.71mg to 4.51 ± 1.6mg within 5 days and 15.44 ± 3.18mg to 5.12 ± 2.04 mg within 3 days, respectively. Whereas, 1% gelatin with 0.5% sodium alginate exhibited a degradation profile, with a reduction in weight of 50%, from 40 ± 2 mg to 20 ± 1.5 mg, over the course of 7 days, which was faster than that of pure alginate hydrogels, highlighting the influence of gelatin on the hydrogel’s stability and degradation dynamics (Figure 5 b). For further studies, 0.5% alginate and 1% gelatin with 0.5% alginate were further used for analysis due to their degradation and swelling profiles.

### Shear stress rate of hydrogels

The rheological behavior of 0.5% sodium alginate and 1% gelatin–0.5% sodium alginate composite hydrogels was assessed using a shear rate sweep ranging from 0.1–100 s⁻¹ at 25°C (n = 2). Both hydrogels exhibited distinct non-Newtonian shear-thinning behavior. The composite hydrogel reached a peak shear stress of ∼75–80 Pa, followed by slight stabilization, indicative of improved structural integrity and resistance under shear forces. In contrast, the alginate hydrogel alone exhibited a sharp peak exceeding 80 Pa at lower shear rates (15–20 s⁻¹), followed by a continuous decline in shear stress, suggesting significant structural breakdown under applied stress (Figure 5d).

The yield stress, determined from the flow curve deviation, ranged from 25 to 50 Pa for the alginate hydrogel, with the composite hydrogel displaying comparatively higher yield stress values. Young’s modulus, calculated from the linear region (0–10% strain) of the stress–strain curve, was 0.08 kPa for alginate and 0.10 kPa for the composite hydrogel, indicating a modest increase in stiffness upon gelatin incorporation.

### Viscosity of hydrogels

The viscosity profiles of the hydrogels were evaluated as a function of shear rate (Figure 5e; Table 1). All formulations exhibited high viscosity at lower shear rates, followed by a pronounced decrease with increasing shear rate, confirming non-Newtonian shear-thinning behavior.

**Table 1:**
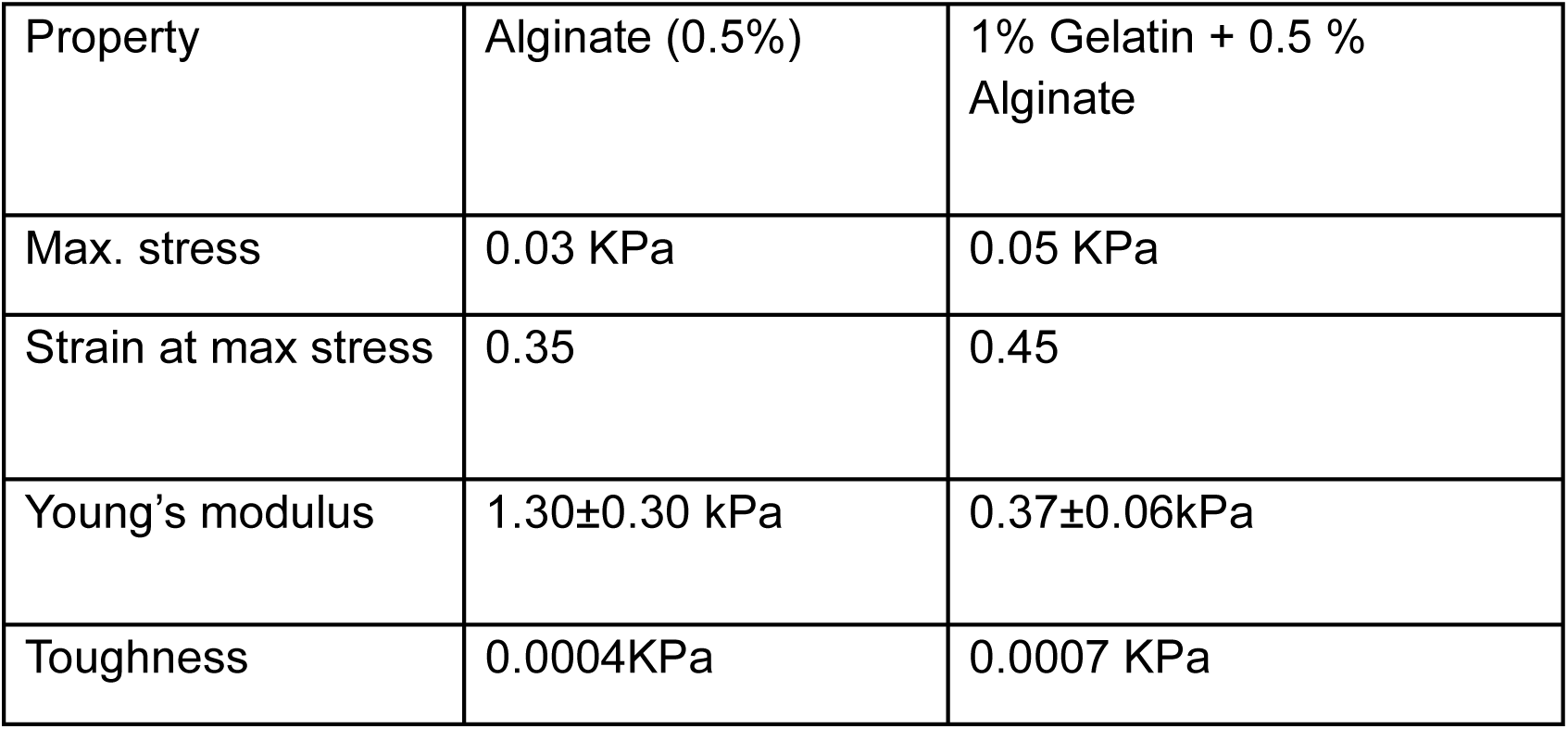
Viscosity analysis of 0.5% Sodium alginate and 1% gelatin with 0.5% sodium alginate.

Among the tested formulations, 0.5% sodium alginate consistently demonstrated the lowest viscosity across the measured shear rate range. The incorporation of dECM increased the viscosity of both alginate alone and gelatin–alginate composite hydrogels, indicating enhanced network interactions within the matrix.

At higher shear rates (∼15 s⁻¹), the viscosities of all hydrogels converged to comparable values ranging between 500 and 1000 mPa·s. The gelatin–alginate composite exhibited a steeper reduction in viscosity than alginate alone, suggesting greater structural rearrangement under shear stress. A steeper viscosity reduction was observed for the gelatin-alginate mixture than for alginate alone.

### Brunauer–Emmett–Teller (BET) analysis

Surface area analysis using the BET method was performed to characterize the pore architecture and surface properties of the hydrogels. The results revealed that alginate to exhibited a higher surface area (BET: 1.996 m²/g; Langmuir: 15.945 m²/g; Figure 6 a-e, Table 2) than the gelatin alginate composite (BET: 0.285 m²/g; Langmuir: 0.539 m²/g) (Table 2). Pore-size distribution analysis indicated that the composite hydrogel exhibited mesoporous characteristics, with pore diameters ranging from 2 to 50 nm. In contrast, alginate alone showed comparatively larger pore diameters of 62–98 nm, suggesting increased pore expansion and structural openness. The higher surface area and larger pore dimensions observed in alginate hydrogels indicate greater porosity than in the composite formulation.

**Figure 6:**
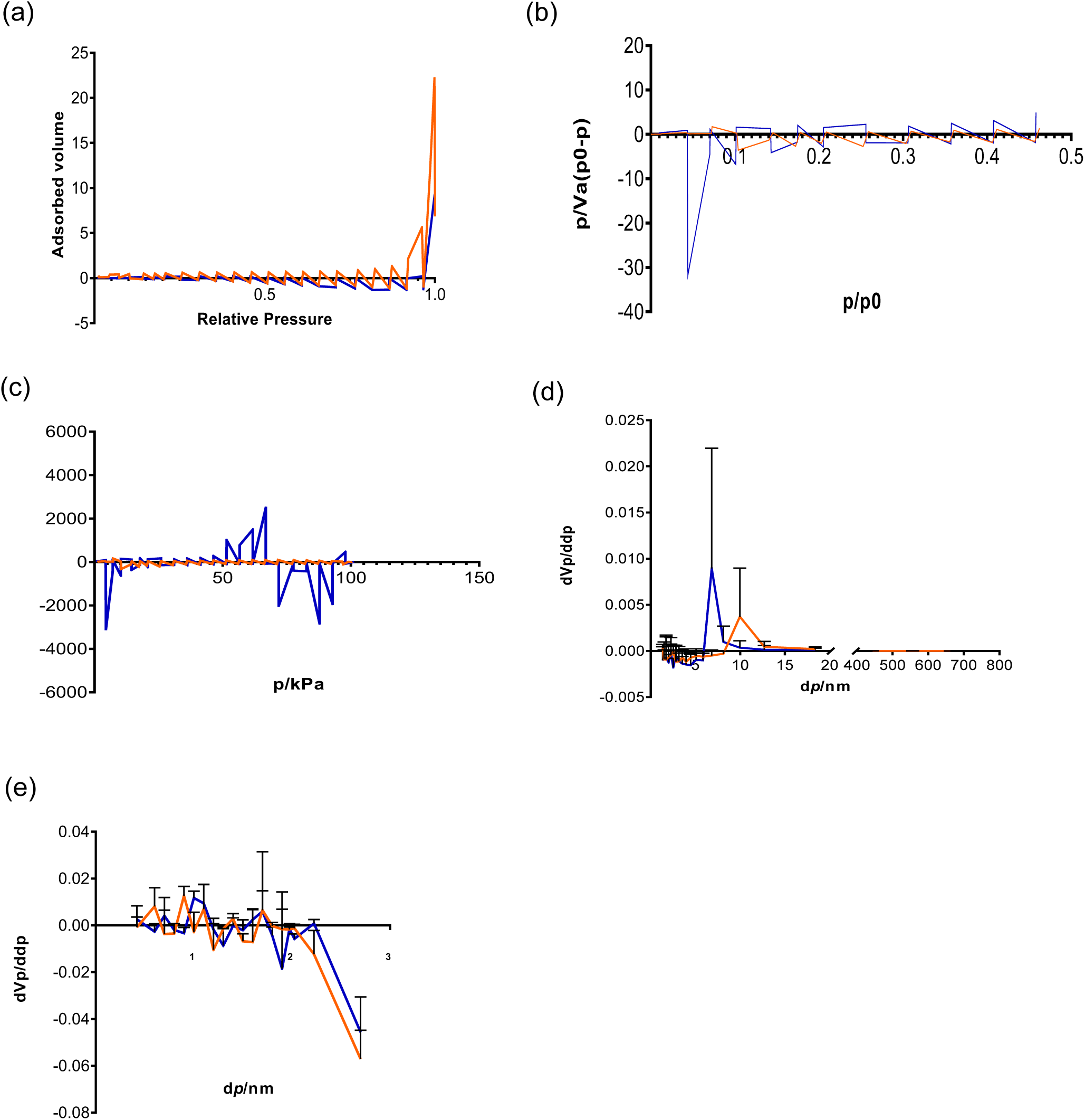
(a-e) BET surface area analysis of the hydrogels: (a)N_2_ -adsorption−desorption isotherm plots, (b) BET surface area, (c) Langmuir surface area, (d) BJH Pore Size Distribution graph for Pore Volume and (e)Micropore graph for average pore diameter of 0.5% sodium alginate and composite hydrogel of 0.5% sodium alginate and 1% gelatin [orange-0.5% sodium alginate; blue-1% gelatin+0.5% sodium alginate]

**Table 2:** BET analysis of 0.5% Sodium alginate and 1% gelatin with 0.5% sodium alginate.

| Hydrogel Type | BET Surface Area (m<sup>2</sup>/g) | Langmuir Surface Area (m<sup>2</sup>/g) | Pore Volume (cm<sup>3</sup>/g) | Avg. Pore Diameter (nm) |
| --- | --- | --- | --- | --- |
| Alginate | 1.996 | 15.945 | 0.026994 | 61.692 |
| Gelatin with sodium alginate | 0.28504 | 0.5394 | 0.00785 | 4.8734 |

### 3D ovary development

The ovarian structure was developed by integrating a medullary layer composed of 1% gelatin and 0.5% alginate, and a cortical layer composed of 0.5% alginate beads mixed with CHO cells (Figure 7 b). A low concentration of extracellular matrix (ECM) proteins (1 µg/mL) was incorporated into both polymer components, and the structure was cultured for 7 days. The synergistic effects of the combined hydrogel components, along with ECM proteins, significantly improved cell proliferation in this zonal strategy (161.03 ± 09, p<0.0001) (Figure 7 a) compared to cells cultured in alginate with dECM alone.

**Figure 7:**
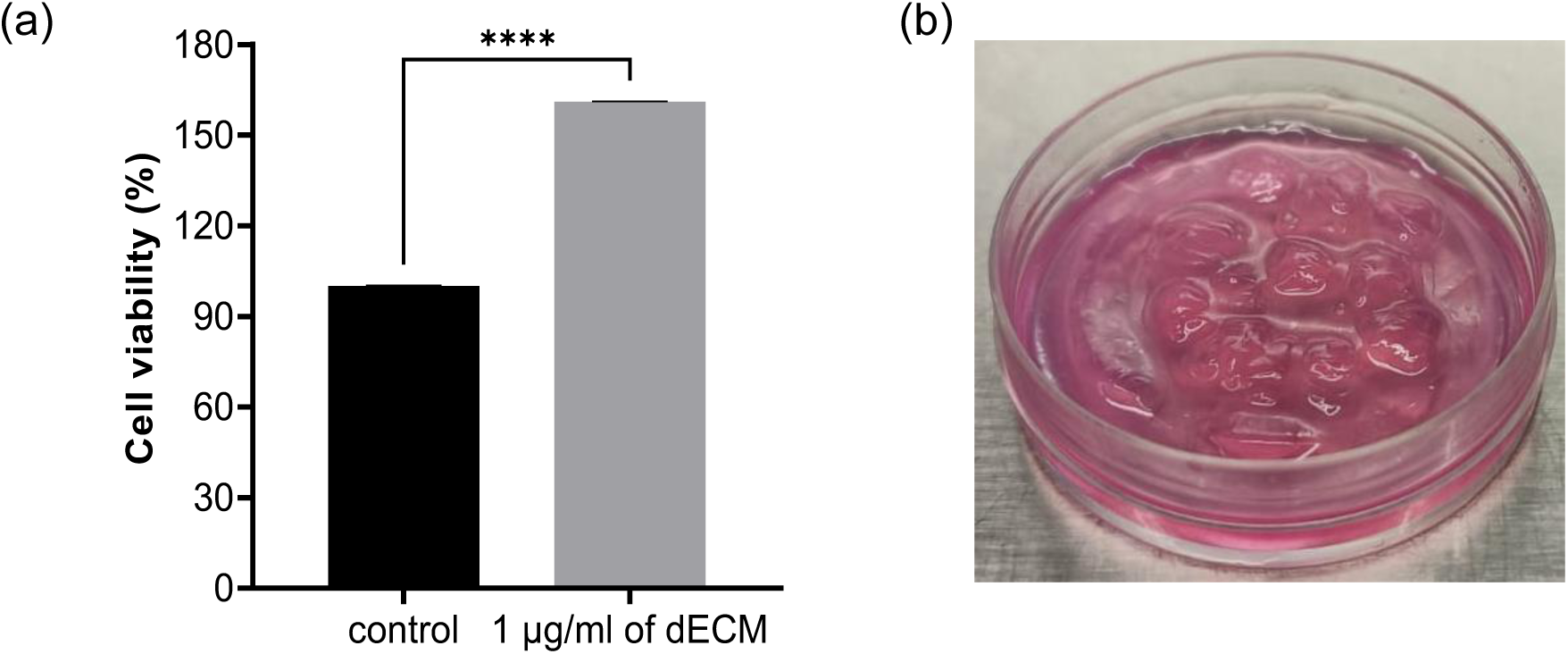
(a) MTT analysis for the 1µg/ml of dECM with 0.5% sodium alginate (p<0.0001 Vs control), (b) Representative images of 3D ovary structure with zonal architecture after 7 days of culture with cells cultured in 0.5% of sodium alginate with 1% gelatin with 0.5% sodium alginate as the base.

### Discussion

This study establishes that zonal hydrogel scaffolds incorporating ovarian dECM improve cell viability and more accurately recapitulate native ovarian mechanics compared to homogeneous systems, advancing the development of functional artificial ovaries. In this study, we engineered a bilayer construct that recapitulated region-specific ovarian mechanics using 0.5% alginate and composite gelatin-alginate hydrogels that mimic the mechanical properties of the cortex and medulla. By integrating ovarian extracellular matrix proteins into these polymers, this dynamic scaffold significantly increased cell proliferation.

The mechanical heterogeneity of ovarian tissue is functionally significant, as the rheological characteristics of the human ovarian cortex differ between reproductive and nonreproductive stages(18). The Young’s modulus of the human ovarian cortex is approximately 3.1 kPa during reproductive years, in contrast to 1–2 kPa for the medulla(35). To mimic the natural ovarian ECM, an intermediate yield stress could balance structural support with cellular dynamics. Recent advancements in the field of ECM biology have utilized dECM in the development of artificial organs, as it preserves ECM proteins, enabling faithful reproduction of the complex *in vivo* environment *in vitro* for potential drug testing and organ reconstruction(36). We observed successful decellularization, with complete removal of nuclear material and maximum retention of proteins, as evidenced by SEM (Figure 3). This is similar to previous studies that have also used both mechanical and chemical techniques for ovarian decellularization in animal species, such as mice, pigs, goats, and sheep(21,37,38). The solubilization of ovarian dECM enabled its homogeneous incorporation within hydrogel matrices, ensuring consistent cell–matrix interactions throughout the construct(39). The enhanced cellular proliferation observed following dECM supplementation (Figure 4) suggests that ovarian ECM components provide essential biochemical cues that are absent in homogenous alginate hydrogels. While pure alginate lacks cell-adhesive ligands, dECM-derived proteins promote integrin-mediated adhesion and activate survival pathways, such as FAK/ERK signalling, which are critical for cellular attachment, proliferation, and function(40). However, dose-dependent toxicity of dECM was observed, especially at higher concentrations of ECM proteins, suggesting the presence of toxic soluble ECM fragments that were not rendered inert(41). The increased stiffness observed in alginate (Figure 5d) and ionic calcium cross-linking could also limit nutrient and oxygen diffusion in the hydrogels, resulting in oxidative and hypoxic stress in the growing cells(42,43). In contrast, the composite gelatin alginate hydrogel was more tolerant toward increased concentrations of ECM proteins (Figure 5e). The presence of RGD motifs in gelatin is known to improve integrin binding, increase porosity, and reduce stiffness in this hydrogel(44). This finding emphasizes the importance of optimizing the dECM concentration to balance biochemical signalling with mechanical properties and diffusion characteristics.

Although dECM alone provides biochemical fidelity, its integration within mechanically defined hydrogels enables simultaneous control over structural integrity and cell–matrix signalling, a critical requirement for functional ovarian reconstruction. The combination of dECM supplementation with the zonal strategy significantly enhanced CHO cell proliferation (Figure 7), which can be attributed to the synergistic effects of appropriate mechanical stiffness and biochemical cues, coupled with improved nutrient diffusion within the bilayer architecture. Previous studies support this finding, as alginate is the most commonly used polymer for *in vitro* culture of follicles and ovarian cells(45,46). In the current study, the mechanical properties of 0.5% alginate were similar to those reported for native human ovaries, with a shear strength of 121 ± 7 Pa, rigidity of 3.2kPa, and Young’s modulus of 0.84 ± 0.16 kPa(47,48). BET analysis revealed the microporous and mesoporous nature of the alginate hydrogel, with an optimal viscosity that can accommodate volume changes during follicle development while maintaining structural integrity. The use of collagen polymers abundant in ovarian ECM, as well as collagen organoids, was found to result in faster degradation, even though they increased follicle viability(49,50). This presents an interesting duality, in which the benefits of increased viability must be balanced with the rate of material degradation in tissue engineering applications.

Despite the promise of homogeneous alginate and collagen polymers used for three-dimensional ovarian constructs, they exhibit limitations. The use of homogenous synthetic polymers, such as poly ethylene glycol (PEG), has been reported; however, its potential for follicle maturation is compromised by the absence of RGD motifs critical for cell adhesion and function(51). The importance of selecting appropriate polymer combinations to optimize biological functions and tissue responsiveness is further emphasized by these studies. Recent studies have identified synergistic effects achieved through the combination of natural polymers, such as gelatin, hyaluronic acid (HA), and collagen, to improve the cellular interactions of alginate(52). In a study, Khunmanee et al.,(53) successfully developed chitosan-co-thiolated hyaluronic (CSHS (0.5%) hydrogel for mouse preantral follicle culture and obtained successful MII oocyte release when compared to 0.5% alginate alone. The G’ was found to be approximately 85 Pa, and this stiffer property was analyzed for improved follicle development in this 3D culture system.

In alignment with these observations, the present study also demonstrated limited cell proliferation of CHO cells in homogeneous 0.5% alginate scaffolds, whereas supplementation with dECM proteins had improved cell proliferation (Figure 4). Jamalzaei et al.,(54) previously cocultured preantral follicles in alginate hydrogels with ovarian cells to increase survival rate. This again supports the hypothesis that protein and growth factor supplementation is required to improve cell proliferation in homogeneous alginate hydrogels. However, in the present study, combining dECM proteins with zonal mechanical gradients significantly improved cellular proliferation, demonstrating that both the biochemical composition and mechanical compartmentalization are critical for functional scaffold design.

To develop artificial ovaries that support the growth of ovarian cells, including follicles, several bioengineering methods have been employed, ranging from simple alginate hydrogel droplets to precise biomimetic bioprinting(55). Although previous studies have successfully grown secondary follicles to antral stages in various animal species using polymer-based scaffolds, the *in vitro* development of human primordial and preantral follicles remains challenging. While the present model recapitulates key mechanical and biochemical aspects of the ovary, future studies incorporating vascular networks and primary human ovarian stromal cells will further enhance physiological relevance. This study employed CHO cells as a proof-of-concept model to assess cellular viability and proliferation within the scaffold. Although CHO cells provided valuable insights into biocompatibility and the effects of mechanical and biochemical cues, they do not recapitulate the complex biology of ovarian cells. Future validation with primary granulosa cells, theca cells, and intact follicles at various developmental stages is essential to confirm biological relevance and assess functional outcomes, including steroidogenesis, oocyte maturation, and follicle growth kinetics. Future studies are also required to incorporate vascularization strategies for vascular network formation, such as co-culture with endothelial cells or the incorporation of pro-angiogenic growth factors within the medullary compartment.

In summary, this study will have a profound influence on the development of physiologically relevant three-dimensional ovarian models for reproductive medicine and biomedical research. Such platforms can serve as *in vitro* models for exploring ovarian function and pathologies, offering substantial potential for drug testing, toxicology screening, and personalized fertility preservation strategies. The zonal biomaterial approach presented herein, utilizing alginate for the cortical region to foster a mechanically stable environment for early-stage follicles and gelatin-alginate for the medullary region to provide biochemical signalling and stromal support, represents a significant advancement toward functional artificial ovaries.

## Supporting information

https://learnermanipal-my.sharepoint.com/:w:/g/personal/nikhila_mcbrmpl2025_learner_manipal_edu/IQBL5yw4EnuSTKPvJ15tDA-DAbAYL6MVErKm4C-HWrk9Vcs

## Ethical Compliance

For obtaining the ovine ovaries, Institutional Animal Ethical Clearance of Kasturba Medical College, MAHE, Manipal was obtained. IAEC No-IAEC/KMC/15/2023.

## Data Access Statement

Not Applicable.

## Author Contributions

NSK and SSM were involved in the development of hydrogel and cytotoxicity analysis. VVM, GK, and SGK contributed to the decellularization and characterization of ovarian tissue. AS, BNS, and KV focused on the mechanical property characterization of hydrogels. RNS was responsible for supervision and editing of the manuscript, while RN handled the conception, manuscript preparation, and editing.

