## Supplementary material for "DEVELOPMENT OF A BIOMIMETIC 3D OVARIAN SCAFFOLD USING DECELLULARIZED EXTRACELLULAR MATRIX AND MECHANICALLY TUNED HYDROGELS": https://learnermanipal-my.sharepoint.com/:w:/g/personal/nikhila_mcbrmpl2025_learner_manipal_edu/IQBL5yw4EnuSTKPvJ15tDA-DAbAYL6MVErKm4C-HWrk9Vcs

Supplementary Figure 1: Effect of ECM proteins on oocyte maturation: a) Graph representing the maturation potential of mouse oocytes cultured in varying concentrations of solubilized dECM, b) Representative images of mouse oocytes post 18 h of incubation, c) Graph representing the ROS level in mouse oocytes cultured in varying concentrations of solubilized dECM, d) Representative images of DCHF-DA stained mouse oocytes.

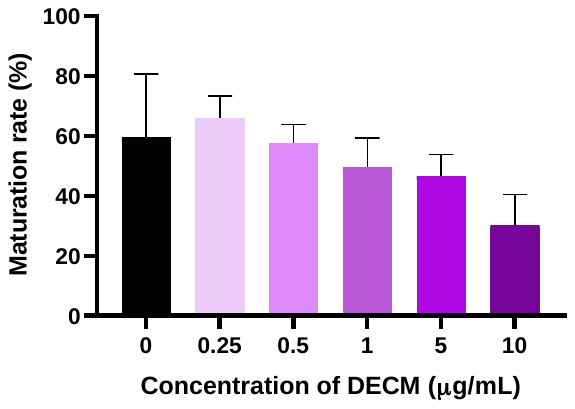

(b)

(a)

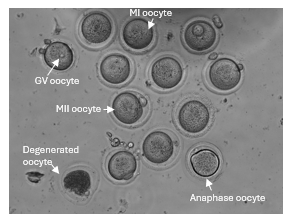

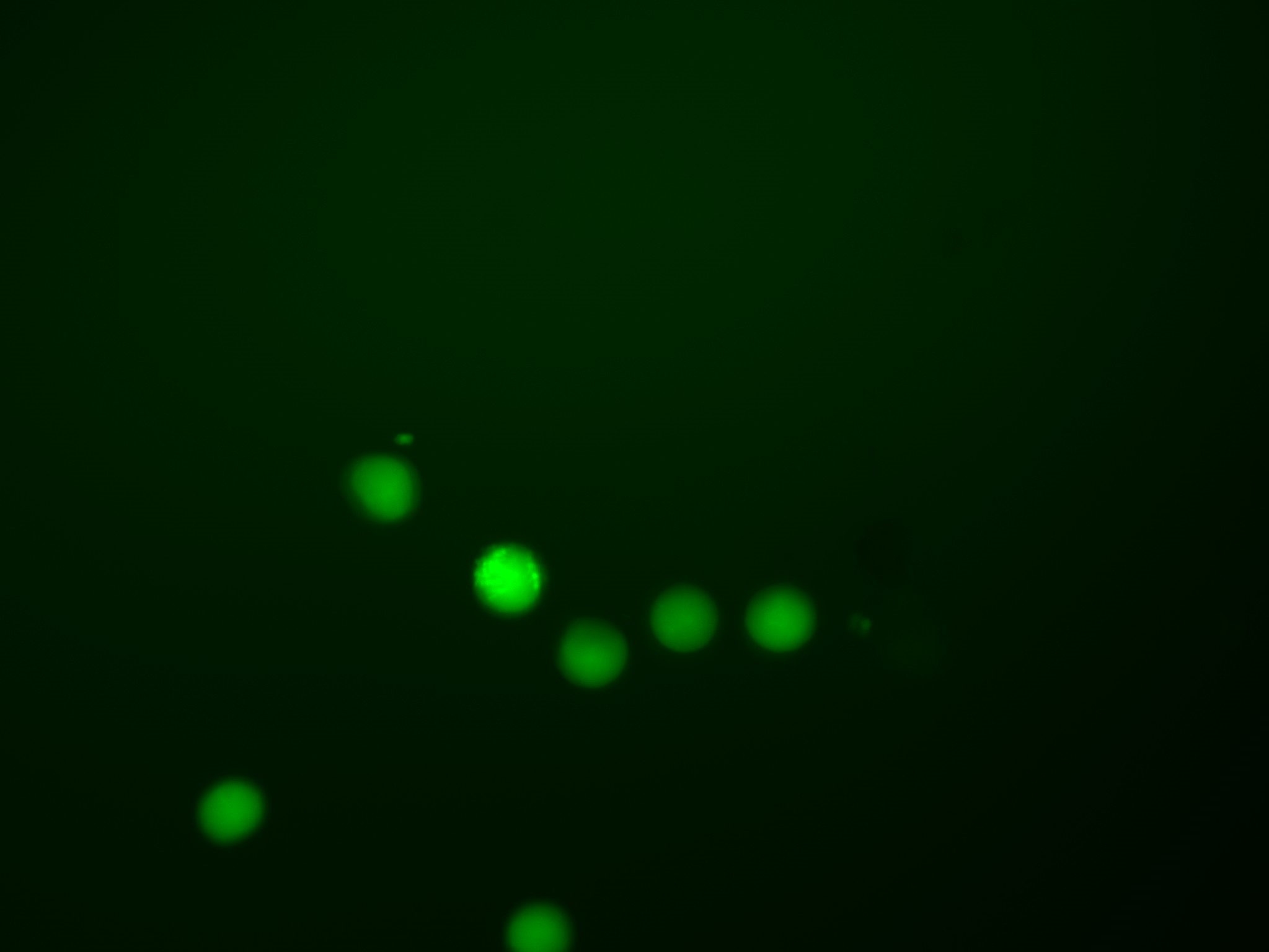

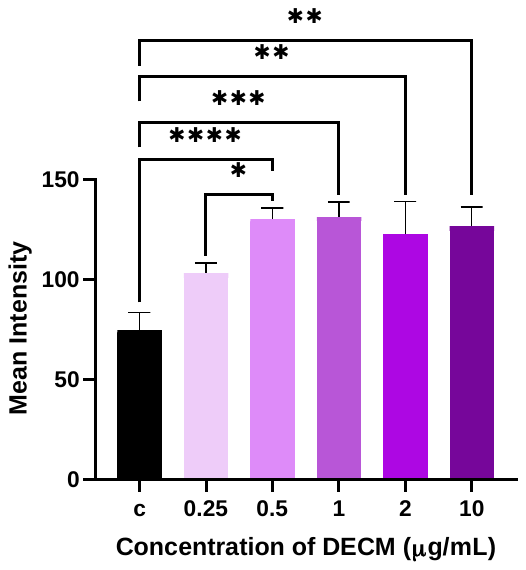

(d)

(c)

Anaphase oocyte
